# Application of synthetic peptide CEP1 increases nutrient uptake rates along plant roots

**DOI:** 10.1101/2021.10.11.463963

**Authors:** Sonali Roy, Marcus Griffiths, Ivone Torres-Jerez, Bailey Sanchez, Elizabeth Antonelli, Divya Jain, Nicholas Krom, Shulan Zhang, Larry M. York, Wolf-Rüdiger Scheible, Michael K. Udvardi

## Abstract

The root system of a plant provides vital functions including resource uptake, storage, and anchorage in soil. Uptake from the soil of macro-nutrients like nitrogen (N), phosphorus (P), potassium (K), and sulphur (S) is critical for plant growth and development. Small signaling peptide (SSP) hormones are best known as potent regulators of plant growth and development with a few also known to have specialized roles in macronutrient utilization. Here we describe a high-throughput screen of SSP effects on root uptake of multiple nutrients. The SSP, MtCEP1 enhanced nitrate uptake rate per unit root length in *Medicago truncatula* plants deprived of N. MtCEP1 and AtCEP1 enhanced uptake not only of nitrate, but also phosphate and sulfate in both Medicago and Arabidopsis. Transcriptome analysis of Medicago roots treated with different MtCEP1 encoded peptide domains revealed that hundreds of genes respond to these peptides, including several nitrate transporters and a sulfate transporter that may mediate the uptake of these macronutrients downstream of CEP1 signaling. Likewise, several putative signaling pathway genes were induced in roots by CEP1 treatment. Thus, a scalable method has been developed for screening synthetic peptides of potential use in agriculture, with CEP1 shown to be one such peptide.

## Introduction

The root system of a plant provides vital functions including resource uptake, storage, and anchorage in soil. For plant growth and development, uptake from the soil of macronutrients, i.e. nitrogen (N), phosphorus (P), potassium (K), and sulphur (S), and micronutrients is critical (Hawkesford and Barraclough, 2011). Soil macronutrients are often present at limiting concentrations for optimal crop yield. Therefore chemical fertilizers are widely used to enrich soils and enhance crop productivity, although their use comes at significant economic and environmental costs (Fageria, 2008). Currently, fertilizer use in agriculture is neither sustainable nor efficient; with as little as 10-30% of applied fertilizer being captured by crop roots (Wortmann,2014), leading to fertilizer losses through leaching, erosion and gaseous emissions, with concomitant eutrophication of inland and marine waters and addition of greenhouse gases to the atmosphere. Hence, understanding the molecular mechanisms governing plant nutrient uptake, which may enable new approaches to increase the efficiency of fertilizer use, is important.

Small signaling peptides (SSPs), also called peptide hormones, are best known for their influence on plant growth and development, with a few peptides also known to influence nutrient uptake and/or assimilation (Matsubayashi, 2014; de Bang et al., 2017a; Roy et al., 2018). The role of SSPs in regulating root system architecture in response to biotic and abiotic factors is of growing interest. Plant genomes may encode thousands of SSPs. For example, 1800 putative SSP genes have been annotated in the legume, *Medicago truncatula*, while >1000 have been identified in *Arabidopsis thaliana* (Ghorbani et al., 2015; de Bang et al., 2017b). SSPs, which result from processing of longer, precursor polypeptides, range in size from between 5-75 amino acids and are perceived by plasma-membrane receptors of the leucine-rich repeat receptor like kinase (LRR-RLK) class (Wang et al., 2020). Peptides are usually encoded in the C-terminal part of the precursor polypeptide and have conserved residues within their sequences that are shared with other SSPs of the same “family” (Tavormina et al., 2015). SSP family members often control similar processes (Murphy et al., 2012). However, variation within families of a given species is also known to exist (Ogilvie et al., 2014). Additionally, sequences are often conserved across species explaining activity of peptides in distantly related species (Oelkers et al., 2008; Hastwell et al., 2017). SSPs can exert their effects locally or systemically because of the ability of some to be transported via the vasculature (Notaguchi and Okamoto, 2015). Chemically synthesized forms of SSP may be recognized by cell-surface receptors thereby retaining their morphogenic properties (Okuda et al., 2009; Imin et al., 2013). Synthetic peptides therefore provide an invaluable tool for researchers to uncover novel functions of plant SSPs within days of peptide treatment. Interestingly, some synthetic peptides with no apparent homology to plant-SSPs can also alter plant development, opening up interesting avenues for the development of novel plant growth and physiology regulators (Bao et al., 2017).

In *Arabidopsis thaliana*, the peptide AtCEP1 (C-TERMINALLY ENCODED PEPTIDE) is induced in roots grown in soils with heterogeneous nitrogen availability (Tabata et al., 2014; Ohkubo et al., 2017). Application of synthetic AtCEP1 induced expression of the nitrate uptake transporters *AtNRT1*.*1/AtNPF6*.*3 (NITRATE TRANSPORTER 1*.*1/NITRATE PEPTIDE TRANSPORTER FAMILY 6*.*3), AtNRT2*.*1* and *AtNRT3*.*1* in roots, indicating a role for AtCEP1 in nitrate absorption (Tabata et al., 2014). Although some studies have shown that SSPs can affect nutrient uptake, few, if any, systematic screens have been undertaken to identify physiological effects of synthetic peptides. Here, we have established a hydroponics-based plant growth system and an effective protocol for measuring the effects of synthetic SSPs on depletion rates from the medium of a range of different nutrients, using ion chromatography. We demonstrate the reliability of this system in measuring higher nitrate uptake rates 48 hours post treatment with the *M. truncatula* orthologue of AtCEP1 peptide, MtCEP1, compared to a no-peptide treatment. Surprisingly, we found that synthetic CEP peptides also enhanced root uptake of phosphate and sulfate. RNAseq analysis showed that MtCEP1 Domain1 peptide had the strongest effect on the Medicago root transcriptome and revealed putative new targets of CEP1 signaling.

## Materials and Methods

### Hydroponic plant growth

#### Medicago truncatula

*M. truncatula* jemalong A17 seeds were scarified, sterilized, plated on water agarose medium and transferred to 4°C for three days in the dark. Seeds were allowed to germinate in the dark at 23°C for 16 hours. Germinated seedlings were transferred to Broughton & Dilworth (B&D) nutrition medium (with 1% Agarose) in ‘filter paper sandwich’ systems and grown under short-day conditions (8-/16-h day/night cycle) with 120 mol m^2^ s^-1^ light for four days (Breakspear et al., 2014). Four day old seedlings were sown in cutouts of Identi-Plug foam (Jaece Industries Inc., NY, USA) and 15 mL falcon tubes with the bottom cone cut away, and placed into aerated hydroponic tanks containing B&D full nutrition medium (Table S1) (Figure 1A). Plants were grown under short-day conditions in Conviron walk-in plant growth rooms at 22 °C temperature and 120 mol m^2^ s^-1^ light for 11 additional days. Prior to the uptake experiment, the plants were then transferred to a macronutrient-free nutrient solution (500 μM CaCl_2_, 1000 μM MES) with the respective peptide treatment of 1 μM and micronutrients unless otherwise stated, for 48 hours before measurement (Figure 1B).

**Figure 1.**
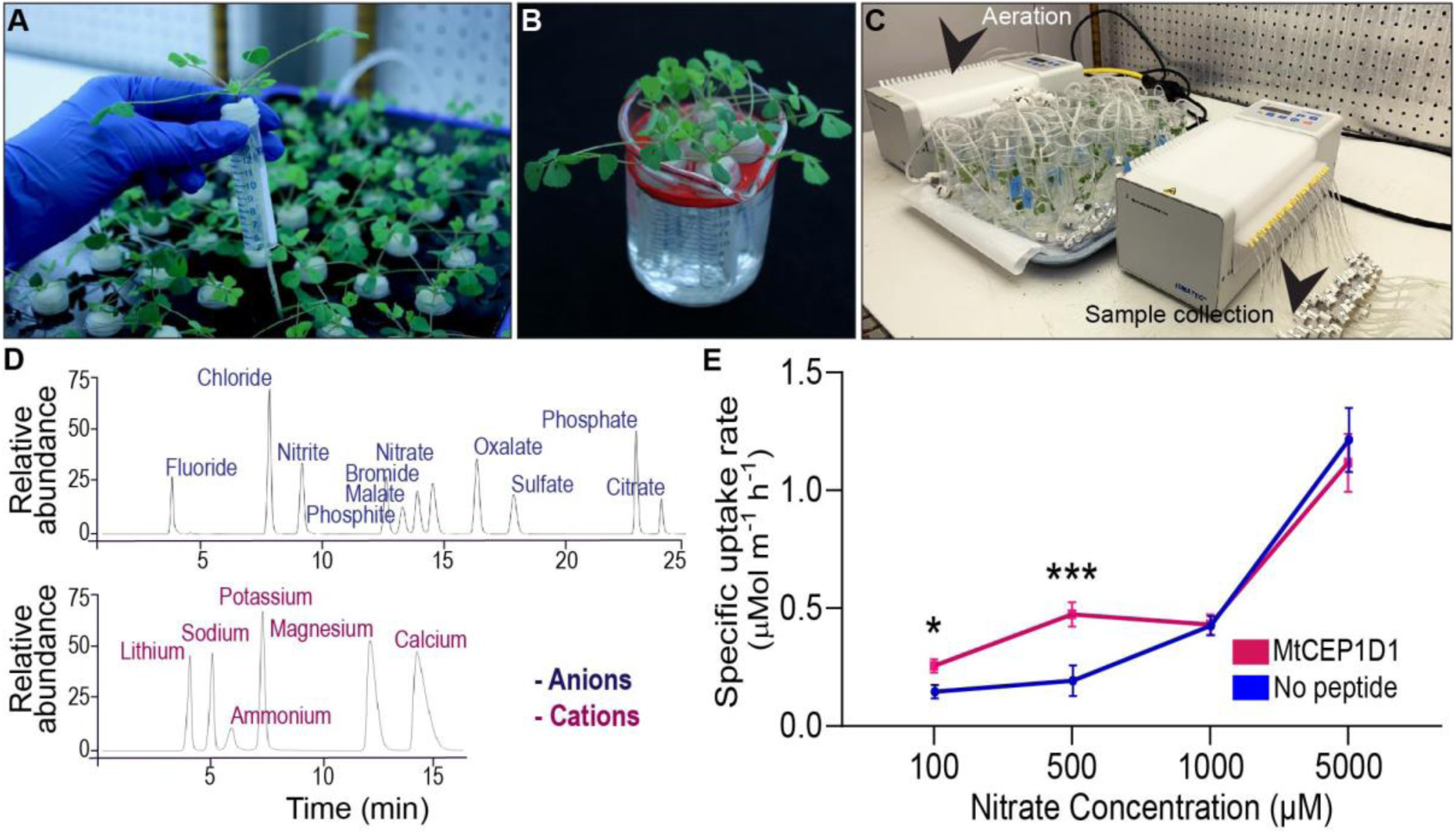
A phenotyping platform for determining uptake rates of multiple ions simultaneously. (A) *Medicago truncatula* plants grown in aerated hydroponic tanks for eleven days. (B) Treatment of plants in nutrient deprivation solution for 48 hours with 1 μM added peptide of interest. (C) Nutrient uptake assay consisting of 24 hydroponic chambers with one plant each. (D) Determination of nutrient concentrations in collected samples by ion chromatography. Time of elution determined for eleven cations and six anions using known standard solutions. (E) Enhanced specific nitrate uptake rate in the high-affinity range (100-500 uM) resulting from pre-treatment with 100 nM MtCEP1D1 Student’s t-test \**p*<0.05, \*\*\**p*<0.001. n=4-6 per sample.

#### Arabidopsis

*A. thaliana* Columbia-0 seeds were surface sterilized (50% Bleach followed by 75% ethanol treatment) and plated aseptically on ½ Murashige & Skoog (with 0.4 % (w/v) Gelzan) and transferred to 4 °C. After two days the seedlings were transferred to 22 °C and grown under short-day conditions (8-/16-h day/night cycle) with 120 mol m^2^ s^-1^ light for four days or till roots were about 1 cm long. These seedlings were then transferred to Falcon 6-well culture plates (Corning, Arizona, USA) containing 5 mL of liquid ½ MS and grown for 10 days on a shaker (80 rpm, New Brunswick platform shaker). Ten plants were pooled together per well to make a biological replicate. The plants were then transferred to a macronutrient-free solution with 1 μM AtCEP1 peptide treatment (Pepscan, the Netherlands), 48 hours prior to the uptake experiment, in order to promote uptake induction.

### Determination of nutrient uptake rates by plant roots using ion chromatography

For the nutrient uptake experiment, plants were processed following the RhizoFlux ions protocol with modifications (Griffiths et al., 2021). A custom ion uptake analysis assay was used with individual plant hydroponic chamber control of nutrient solutions or treatments. The setup consisted of 24 chambers coupled to two peristaltic pumps for nutrient sampling and aeration (Ismatec ISM944A, Cole-Parmer Instrument Company LLC., IL, USA) (Figure 1C). For the Arabidopsis experiments the plants were grown on a shaker and nutrient sampling was conducted with a pipette. Each chamber was filled with a procedure solution containing the respective peptide treatment and macronutrients: (in μM) 100 KNO_3_, 100 NH_4_Cl, 12.5 Ca(H_2_PO_4_)_2_H_2_O, 25 MgSO_4_, 487.5 CaCl_2_, 1000 MES (adjusted to pH6.8 using NaOH). The procedure solution volume used in the Medicago experiments was 35 mL per plant and for the Arabidopsis experiments 15 mL per pool was used. Two minutes after the macronutrient-starved plants were transferred to the individual chambers the first 650 μL nutrient sample was collected. Nutrient solution samples were taken between 0 and 4 h on a deep-well collection plate and the plate was transferred to 4 °C for short term storage and if necessary to -20 °C for long term storage. After the nutrient uptake experiment, the plants were immediately transferred to 4 °C in plastic bags for later root image processing. Ion concentrations of the collected nutrient solution samples were determined using a Thermo Scientific ICS-5000+ ion chromatographic system (Thermo Fisher Scientific, MA, USA) and the data processed to give nutrient concentrations using the Chromeleon 7.2 SR4 software (Thermo Fisher Scientific, MA, USA) (Figure 1D).

### RNA extraction and quantitative PCR

To investigate nutrient responsive effects in *M. truncatula* Jemalong A17, plants were first germinated and grown on full nutrient plates. Four day old seedlings were then transferred to low nitrate (50 μM NH4NO3), low P (6 μM KH2PO4) and sulfate free B & D media (Table S2). After 48 hours, root material from 20 seedlings per biological replicate, was harvested and immediately frozen in liquid nitrogen. For RNA sequencing, three day old Medicago truncatula seedlings grown on water agarose (Life Technologies) medium were treated with 1 μM MtCEP1D1, MtCEP1D2 and AtCEP1 peptide concentrations in water for three hours. For all three biological replicates 20-30 seedling roots were used

Trizol reagent was used to extract total RNA (Life Technologies) following the manufacturer’s protocol (Invitrogen GmbH, Karlsruhe, Germany). Total DNA was digested with RNase free DNase1 (Ambion Inc., Houston, TX) and column purified with RNeasy MinElute CleanUp Kit (Qiagen). RNA was quantified using a Nanodrop Spectrophotometer ND-100 (NanoDrop Technologies, Wilington, DE). RNA integrity was assessed on an Agilent 2100 BioAnalyser and RNA 6000 Nano Chips (Agilent Technologies, Waldbronn, Germany). First-strand complementary DNA was synthesized by priming with oligo-dT_20_ (Qiagen, Hilden, Germany), using Superscript Reverse Transcriptase III (Invitrogen GmbH, Karlsruhe, Germany) following manufacturer’s protocol. Primers were designed using Primer Express V3.0 software. qPCR reactions were carried out in QuantStudio7 (ThermoFisher Scientific Inc.). Five microliters reactions were performed in an optical 384-well plate containing 2.5 μL SYBR Green Power Master Mix reagent (Applied Biosystems), 15 ng cDNA and 200 nM of each forward and reverse gene-specific primer. Transcript levels were normalized using the geometric mean of two housekeeping genes, *MtUBI* (Medtr3g091400) and *MtPTB* (Medtr3g090960). Three biological replicates were included and displayed as relative expression values. Primer sequences are provided in **Supplementary Table 4**.

### RNA-Seq and gene expression analyses

One microgram of total RNA was used to generate RNA-seq libraries using TruSeq Stranded mRNA Library Prep Kit (Illumina Inc.) according to the manufacturer’s protocol. Prior to library construction, RNA integrity and quality were assessed with TapeStation 4200 (Agilent) and only an RNA integrity number (RIN) above nine was used. Size distribution of RNA-seq libraries was analyzed using TapeStation and the libraries were quantified using the Qubit 2.0 Fluorometer (ThermoFisher Scientific) before being shipped to Novogene Inc. for sequencing at 150 bp paired-end with an Illumina Hiseq2000 (Illumina). Data are available on NCBI under SRA number PRJNA764762.

### Root architecture phenotyping

Roots were imaged using a flatbed scanner equipped with a transparency unit (Epson Expression 12000XL, Epson America Inc, CA, USA). The roots were cut away from the shoots, spread out on a transparent plexiglass tray (420 mm x 300 mm) with a 5 mm layer of water (400 mL), imaged in grayscale at a resolution of 600 dpi, and the total root length for each image was analyzed using RhizoVision Explorer v2.0.1 (Seethepalli et al., 2021). One root scan was performed per biological replicate for the uptake experiments, with one scan per Medicago plant, and one scan per pool of Arabidopsis plants.

### Statistical analyses and data evaluation

For the nutrient uptake rate study, data processing to determine specific nutrient uptake rates was conducted using R version 3.6.0 (Team, 2020)(R Development Core Team, 2020) with minor modification to the R code available at https://doi.org/10.5281/zenodo.3893945 (Griffiths *et al*., 2021). The net nutrient depletion and therefore uptake rate by roots was calculated by *In* = (*C*_*t*_ - *C*_0_) / (*t*_0_ - t) where *In* is the net influx into the plant; *C*_0_ is the initial concentration of the solution at the start of the experiment *t*_0_; *C*_*t*_ is the concentration at sampling time. The net uptake rate was then divided by the root system length (cm) to calculate the net specific nutrient uptake rate with the units μmol cm^-1^ h^-1^. Statistical tests were conducted using Graphpad V. 8. The bar in the box plots represents the median values, with each box representing the upper and lower quartiles, and the whiskers representing the minimum and maximum values.

### RNA-Seq mapping and hierarchical clustering

Low quality bases and primer/adapter sequences were removed for quality trimming of each sample using Trimmomatic version 0.36 (http://www.usadellab.org/cms/?page=trimmomatic). Reads less than 30 bases long after trimming were discarded, along with their mate pair. Using HISAT2 version 2.0.5 (https://daehwankimlab.github.io/hisat2/) default mapping parameters and 24 threads, trimmed reads were mapped to an in-house mapped to an in-house re-annotated version of the *M. truncatula* genome release 4.0_reanno (http://bioinfo3.noble.org/). Transcripts were assembled and quantified using Stringtie 1.2.4 (http://www.ccb.jhu.edu/software/stringtie/) with the default assembly parameters. The transcripts identified in control (no peptide) and CEPp treated samples were unified into a single set of transcripts and compared with the reference gene annotation set using Stringtie’s ‘merge’ mode. Differential expression testing was performed using DESeq2 (https://bioconductor.org/packages/release/bioc/html/DESeq2.html). Fold changes were calculated based on average FPKM values and DEG’s selected at a *p*-value cutoff of 0.05 and below.

### Differential gene expression analysis

The threshold for determining the differentially expressed genes (DEGs) was set to a fold change of 1.5 (log2 fold change >|0.58|) and p-value cutoff <0.05. For assessing the common DEGs, all the up and down-regulated DEGs along with shared DEGs between AtCEP1, MtCEP1D1 and MtCEP1D2 were analyzed as Venn diagrams using Venny 2.1.0 (Oliveros, 2016). The up and down-regulated DEGs were enriched for Gene Ontology (GO) terms using the online gene discovery platform, Legume IP V3 (Dai et al., 2021). GO term enrichment tool on the platform extracts GO terms from functional descriptions of protein in UniProt and InterproScan annotations. An adjusted p-value of p<0.05 was used as a cutoff for GO terms to be considered enriched. Unique genes in the top 20 significantly enriched GO terms for all up and down-regulated DEGs were plotted. The up and down-regulated differential expression of known nitrate, phosphate and sulphate transporters were plotted as a heatmap Log2FC>|0.58|), p-value<0.05. From the differentially expressed genes, upregulated kinases and transcription factors were plotted as a heatmap Log2FC>2.0 (corresponding to a 4-fold change in expression level), *p*-value<0.05. All plots were generated using GraphPad Prism 9.0.0 (GraphPad Software, Inc.) and modified using Adobe Illustrator (Adobe Inc. 2021).

## Results

### A platform to measure uptake rates of multiple nutrients reveals that exogenous application of synthetic peptide can directly affect nutrient uptake rates

A new platform for evaluating the effect of synthetic peptides on root uptake of multiple nutrient ions was developed (Figure 1). Plants were grown in a hydroponic system (Figure 1A) and peptides of interest were applied to the nutrient solution around the root system 48 hours prior to nutrient uptake assays (Figure 1B). Plants were then transferred to small assay tubes containing nutrient solution with defined levels of nutrient ions and the respective peptide, which was sampled over a short-duration into a deep-well collection plate (Figure 1C). The anion and cation concentrations of the collected samples were determined using ion chromatography. A decline in ion concentration in the assay solution over time indicated a linear rate of net uptake of the ions (Figure 1D, Figure S1). “Specific” nutrient uptake rates were calculated by dividing ion uptake rate by total root length obtained from image analysis. In a proof-of-concept experiment, exogenous application of Medicago SSP MtCEP1D1 increased the specific rate of nitrate uptake by 70-140% at low external concentrations (100 and 500 μM, p<0.05 and p<0.001, respectively) but not higher concentrations (1 and 5 mM) in treated plants compared to non-treated controls (Figure 1E).

### CEP1 peptide alters root system architecture

To determine if expression of MtCEP1 was regulated by multiple nutrient stresses, we grew *M. truncatula* seedlings on agarose plates containing nutrients for optimal growth (B & D Full Nutrition medium) for three days. Seedlings were deprived of specific macronutrients for 48 hours before quantitative RT-PCR estimation of endogenous *MtCEP1* transcript abundance. Notably, nitrogen deprivation significantly enhanced *MtCEP1* transcript abundance (p<0.01) but not phosphate and sulfate deprivation (Figure 2A). To elucidate the functions of CEP1 peptides and identify key peptide domains, Arabidopsis CEP1 peptide (AtCEP1) and the *M. truncatula* CEP1 (MtCEP1) domain 1 and domain 2 peptides (Figure 2) were applied to agar upon which seedlings were grown, after which root system architecture was analyzed. In agarose, exogenous application of either Arabidopsis or Medicago CEP1 domain 1 (AtCEP1 and MtCEP1D1) significantly reduced lateral root number by ∼50% in Medicago as was previously reported (p<0.01 and p<0.05 respectively; Figure 2BC; Imin et al., 2013). The median number of lateral roots in the presence of MtCEP1D2 was lower than control but was not statistically significant. This effect on root architecture was not observed under conditions used for measuring uptake post 48 hour peptide application (Figure S2).

**Figure 2.**
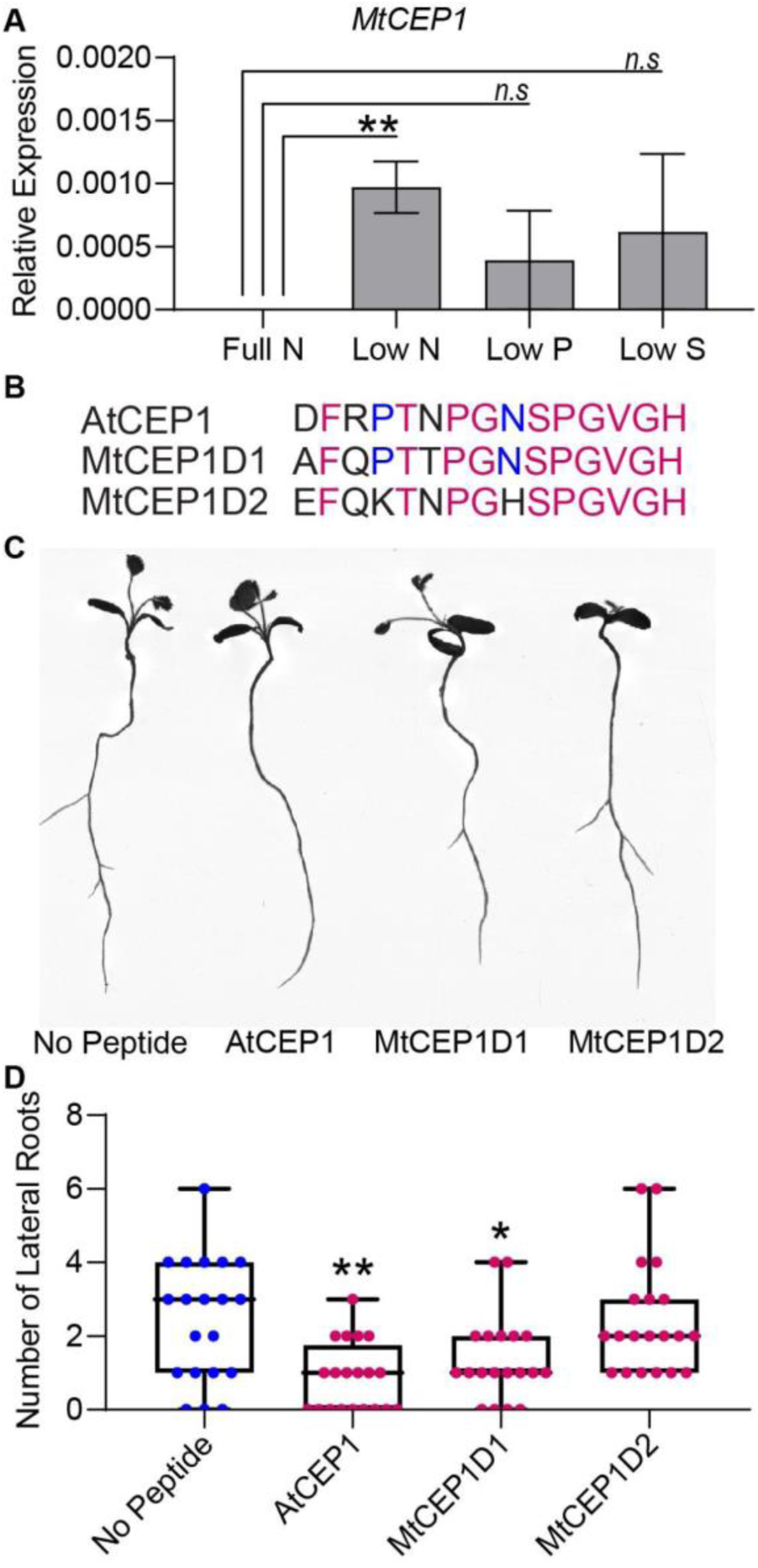
Macronutrient stress responsive expression of CEP1 and effects of synthetic peptides on root system architecture in *Medicago truncatula*. (A) Relative *MtCEP1* transcript abundance in *M. truncatula* seedling roots deprived of a specific macronutrient for 48 hours. Transcript levels were measured by qRT-PCR, normalized to two housekeeping genes, *UBC* and *PTB*, and expressed relative to the level of *MtCEP1* transcript at full nutrition (Full N). Data are averages of three biological replicates in each case. Error bars represent SEM. Student’s t-test \**p*<0.05. (B) Sequences of peptides used in this study. Magenta indicates amino acid residues conserved between all three sequences and blue represents residues conserved between AtCEP1 and MtCEP1D1. Prolines in the fourth and eleventh positions of each peptide were hydroxylated. (C) Representative root scans showing change in root architecture of *M. truncatula* Jemalong A17 seedlings treated with 1 μM peptide compared to no peptide controls. (D) Effect of 1 μM peptide application on lateral root number in *M. truncatula* Jemalong A17 seedlings seven days post germination. One way ANOVA followed by Dunnett’s Multiple comparison test \**p*<0.05, \*\**p*<0.01.

### CEP1 peptides enhance uptake of nitrate, phosphate and sulfate in Arabidopsis and Medicago

The ion uptake platform was used to measure root uptake rates of multiple nutrients simultaneously. In addition to enhancing nitrate uptake rates (*p*<0.01), application of 1 μM of the Arabidopsis AtCEP1 peptide significantly enhanced phosphate and sulfate uptake in *Arabidopsis thaliana* (*p*<0.05; Figure 3A). For *M. truncatula*, both AtCEP1 and Medicago MtCEP1 domain 1 peptide significantly enhanced the nitrate uptake rate (*p*<0.05 and *p*<0.001, respectively; Figure 3B). AtCEP1 peptide did not enhance phosphate or sulfate uptake in *Medicago truncatula*, unlike MtCEP1 that enhanced uptake of both phosphate and sulfate (*p*<0.001 and *p*<0.1, respectively; Figure 3B). Thus, MtCEP1 had a greater effect than AtCEP1 on uptake rates of nitrate, phosphate and sulfate in Medicago (Figure 3B). In contrast, the CEP1 peptides had no effect on ammonium or potassium uptake rates in *Medicago truncatula* (Figure S3).

**Figure 3.**
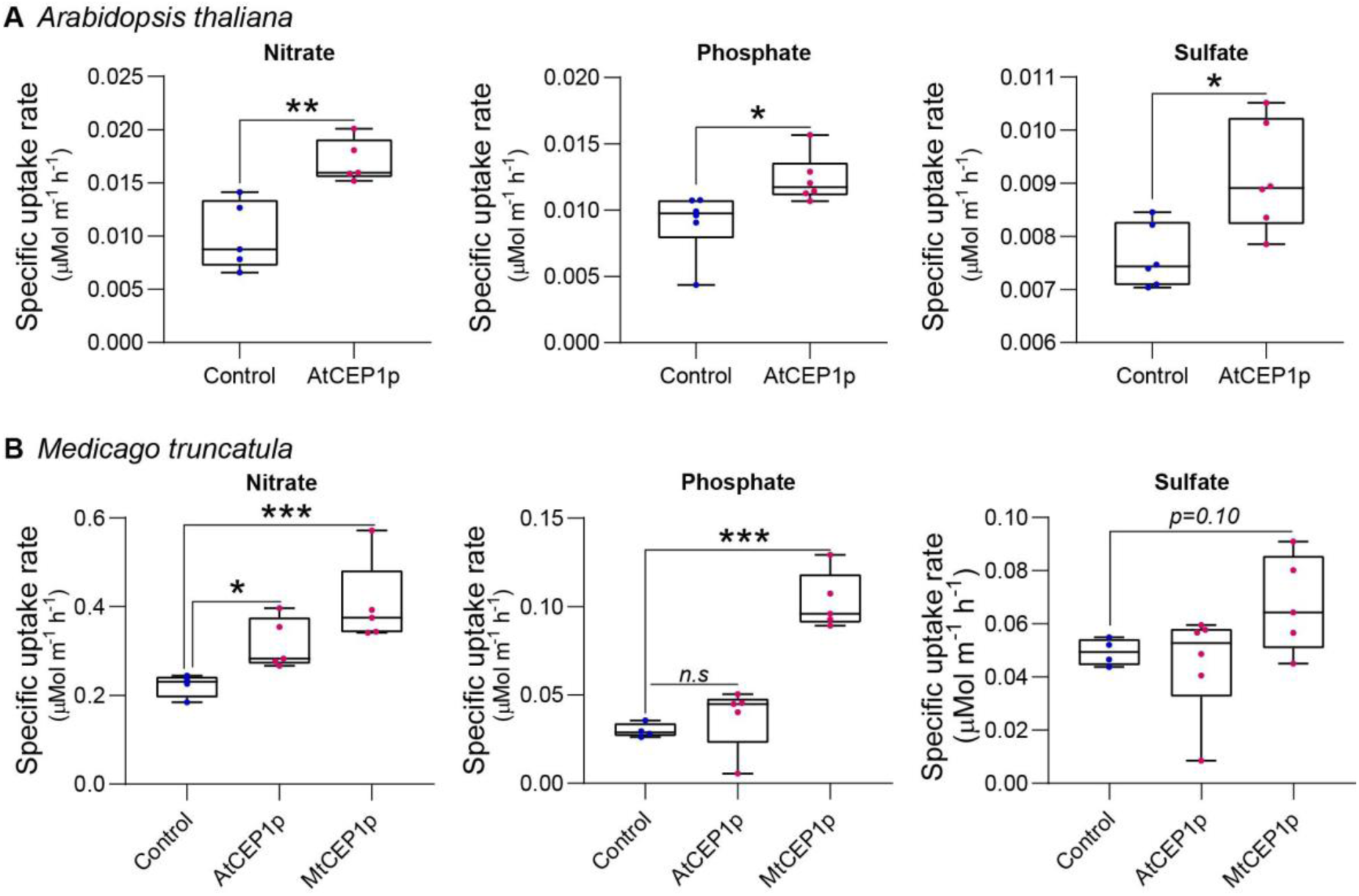
Synthetic CEP1 peptides enhance uptake of nitrate, phosphate and sulfate in *Arabidopsis thaliana* and *Medicago truncatula*. (A) Specific nutrient uptake rates of nitrate, phosphate and sulfate in *Arabidopsis thaliana* in the presence or absence of the synthetic AtCEP1 peptide at a concentration of 1 μM. (B) Uptake rate of nitrate, phosphate and sulfate in *Medicago truncatula* in the presence of synthetic AtCEP1 and MtCEP1 peptide domain 1 at a concentration of 1 μM. Student’s t-test \**p*<0.05, \*\**p*<0.01, \*\*\**p*<0.001. *n*=5-6 per treatment.

### Transcriptome responses of Medicago roots to MtCEP1 peptides

Differential gene expression analysis of RNAseq data from *M. truncatula* seedling roots treated with the three CEP1 peptide variants revealed that MtCEP1D1 triggered the greatest changes in gene expression, with 2,466 genes affected by MtCEP1D1 application of which 1,349 were induced and 1,117 were repressed. Application of CEP1D2 resulted in induction of 1278 genes and repression of 871 genes. Fewer genes in Medicago were affected by treatment of plants with AtCEP1, with only 617 and 482 genes up and down regulated, respectively(Supplementary Table 1). Interestingly, 322 genes were up-regulated and 116 were down-regulated by all three peptide treatments.

Gene ontology enrichment analysis, using Legume IPV3, revealed several biological processes that were affected by CEP1 peptide treatments. Both MtCEP1D1 and AtCEP1 induced expression of genes involved in cell division (GO:0008283 GO:0022402 GO:1903047 GO:0000278 GO:0000280) and cell proliferation (GO:0008284). Both MtCEP1 peptides enhanced the expression of genes involved in pollination (GO:0009856), pollen-pistil interaction (GO:0048544 GO:0009875) and cell recognition (GO:0008037). Genes related to photosynthesis (GO:0015979 GO:0019684 GO:0009765 GO:0009768 GO:0009416 GO:0009767 GO:0009773 GO:0010109) and oxidative stress (GO:0000302 GO:0006979 GO:1901700 GO:0045454 GO:0009651 GO:0006970 GO:0042744 GO:0042743) were down-regulated in response to all three peptides. Genes involved in hormone responses were also affected by CEP1 peptides, with auxin response genes being repressed by both MtCEP1 peptide domains. Upregulation of genes responsible for phosphatase activity (GO:001092) and ABA response (GO:0009738 GO:0071215 GO:0009737) following CEP1D2 application were also found (Supplementary Table 2).

Given the observed increase in nitrate, phosphate and sulfate uptake in response to CEP1 peptides, we looked for changes in the expression of gene families involved in these processes, namely the NITRATE/PEPTIDE TRANSPORTER (NRT/PTR), PHOSPHATE TRANSPORTER (PHT) and SULFATE TRANSPORTER (SULTR) families (Supplementary Table 3). Of the 117 NRT transporter encoding-genes analyzed, 17 were differentially expressed following application of peptides of which seven were induced by MtCEP1D1. One putative sulfate transporter gene, an ortholog of *AtSULTR3;5* (Medtr6g086170), was highly induced by all CEP peptide domains. In our data, we found no PHT phosphate transporter genes significantly induced by the CEP1p application..

Finally, we wanted to identify signaling pathway genes that responded to CEP1 application, which might be interesting targets for breeding crops with enhanced sensitivity to such peptides. We analyzed the top 15% of genes induced by the peptides and focused on those involved in perception and/or relay of signals, especially kinases and transcription factors (Supplementary Table 4). Four kinases: Cyclin dependent Kinase (Medtr8g461270), Serine/Threonine Kinase (Medtr7g056617) and an LRR receptor like kinase (MT4Noble_051661; Medtr2g019170) were highly induced by all three CEP1 peptides. Four Myb transcription factors were also induced by all three peptides. MADS-box transcription factor gene Medtr6g015975 was highly induced by both MtCEP1 domain 1 and 2 peptides, while Medtr4g109830 was induced only by MtCEP1_D2p.

## Discussion

Small signaling peptides are known to perform a wide variety of roles in plant growth and development. However, studies exploiting synthetic SSPs to address agronomically important physiological traits such as root nutrient uptake are scarce. Here, we devised a novel hydroponics-based nutrient uptake screen for high-throughput assessment of SSPs function in modifying root nutrient uptake in Medicago and Arabidopsis. We showed that exogenous application of synthetic SSPs can affect plant nutrient uptake rates, expressed per unit root length to avoid potential confounding effects related to changes in root system architecture. Although treating *M. truncatula* plants with CEP1 peptides for short periods had no effect on total root length. As thousands of SSPs are produced by plants, this nutrient uptake phenotyping screen promises to be valuable for identifying and characterizing novel peptides involved in plant nutrition, which may find application as natural plant growth stimulants in agriculture.

Nitrate is a key macronutrient for plant growth and development and CEP1 peptides play a major role in ensuring plants have sufficient nitrogen for growth when N-availability in soil is heterogeneous or scarce (Tabata et al., 2014; Ohkubo et al., 2017; Laffont et al., 2020). Under N-deficiency stress, roots produce CEP peptides, which serve as ‘N-hunger signals’ that are perceived by receptors in the shoot, which in turn activate further signaling that induces expression of nitrate transporters in roots within N-rich soil patches (Chapman et al., 2020)Tabata et al 2014; (Chapman et al., 2020); Figure 2A). Using the Arabidopsis *cepr* receptor mutants, Tabata et al. (2014), showed that less radiolabelled nitrate accumulated in mutant roots compared to the wild type. Multiple studies demonstrate the effect of externally applied synthetic CEP peptides on root architecture however, effects on nitrate uptake of direct CEP peptide application to plant roots has not been demonstrated before (Imin et al., 2013; Chapman K., et al., 2020). Using a novel nutrient uptake platform, we showed that 48 hour exposure of roots to exogenously applied CEP1 peptides at concentrations of 100 nM and 1 uM can enhance nutrient uptake rates per unit root length of Medicago 70 and 140%, respectively (Figure 1 E). These are physiologically relevant concentrations of nitrate typically found in agricultural soils, which are accessed by so-called high affinity nitrate transporters (Lark *et al*., 2004; Miller *et al*., 2007). At higher nitrate concentrations (>1000 μM), where low-affinity nitrate transport systems dominate, no significant difference was observed between peptide treated plants and controls, indicating that CEP1 peptides control high-but not low-affinity transport of nitrate (Figure 1E). Accordingly, transcriptome analyses revealed that at least seven putative NRT/NPF transporters family genes encoding members of both NRT1 dual affinity transporters and NRT2 high-affinity transporters upregulated by peptide treatment as early as three hours post application (Figure 4C). Since gene overexpression studies fail to discriminate between D1 and D2 peptide domains within the polypeptide sequence encoded by the MtCEP1 gene, our study demonstrates that MtCEP1Domain1 alone is sufficient to induce uptake of nitrate from the surrounding media.

**Figure 4.**
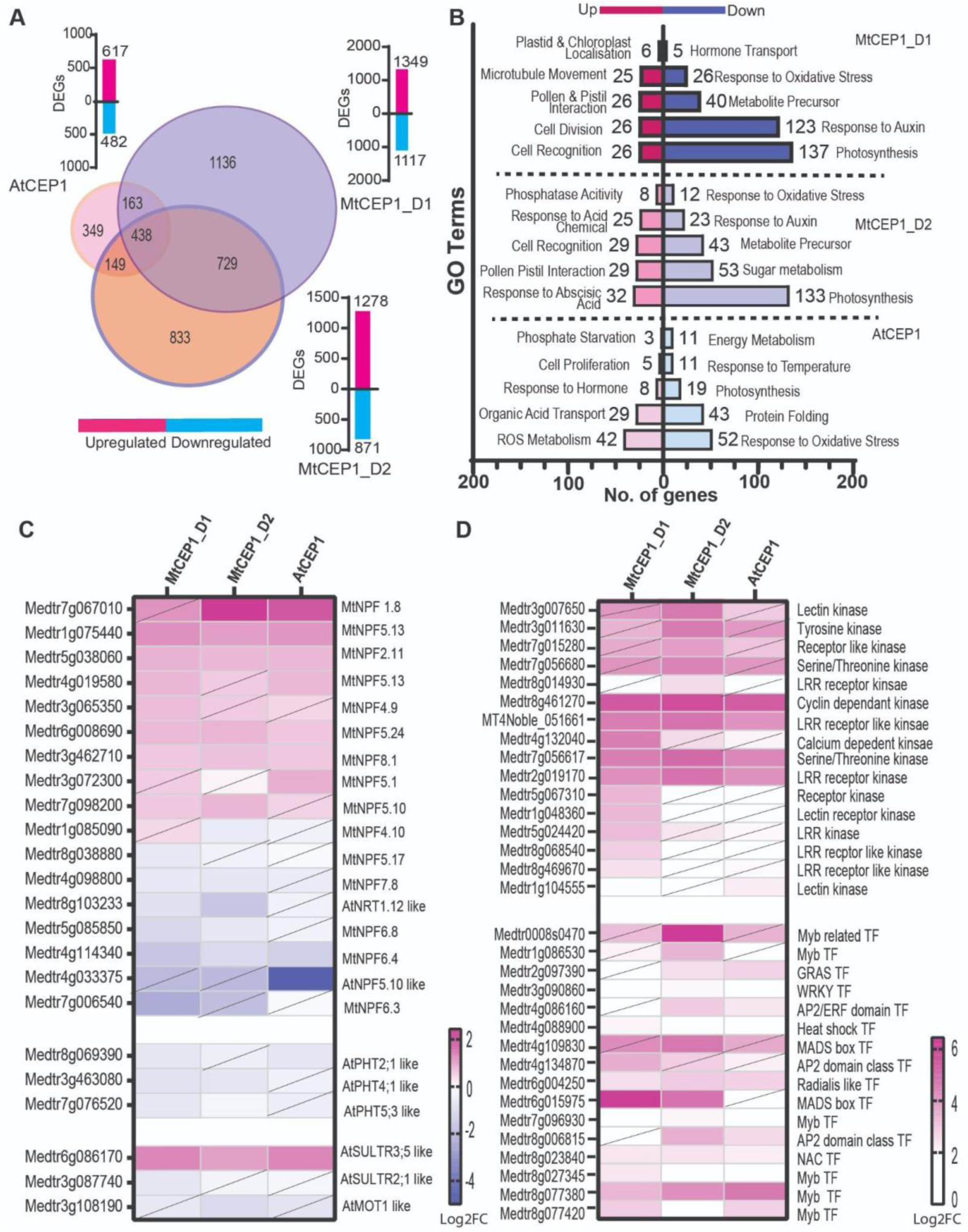
Differential gene expression analysis. (A) Venn diagram showing number of differentially expressed genes following application of AtCEP1p and MtCEP1 peptide domain 1 and 2 in *M. truncatula* (FC>1.5, *p*<0.05). Shared genes are indicated in the overlapping region between peptide treatments. Corresponding histogram shows the total number of DEGs. (B) Histogram showing the top twenty significantly enriched GO terms in up and down-regulated genes (q<0.05). (C) Heat map of putative nitrate and sulphate transporter genes affected by MtCEP1D1, MtCEP1D2 and AtCEP1 peptide treatment in *M. truncatula* (FC>1.5, p<0.05, diagonal line indicates statistically insignificant value). (D) Comparative analysis of CEP1-responsive kinases and transcription factor genes in *M. truncatula* (FC>1.5, p<0.05, diagonal line indicates statistically insignificant value). Average values of three biological replicates are represented. TF stands for Transcription Factor.

Our nutrient uptake methodology can be scaled up or down depending on the seedling size. Using both Arabidopsis (10 plants per replicate) and Medicago (one plant per replicate) we were able to detect measurable changes in uptake of nitrate, phosphate and sulfate within 4-8 hours (Figure 3 A, B, Supplementary Figure 2). Additionally, both the Arabidopsis and Medicago CEP1 domains, AtCEP1 and MtCEP1D1, enhanced Medicago nitrate uptake rate indicating that the CEP signaling pathway and peptide function is conserved across species. Since N-uptake rates induced by MtCEP1D1 were 30% higher than those induced by AtCEP1, some specificity at the species appears to exist, possibly at the level of the peptide receptor which would be expected to have a higher affinity to its endogenous peptide ligand than to that of another plant species . However, given that effects on root system architecture, including important foraging traits such as initiation of lateral roots, are more negatively affected by AtCEP1 than by MtCEP1 peptide domain 1 or 2 (Figure 2 B, C), more work is needed to understand the differential effects of of these peptides. Likewise, further investigation of CEP peptide dosage and length of exposure is required before use in agriculture.

Interestingly, we observed that application of CEP1 on both Medicago and Arabidopsis enhanced uptake not only of nitrate, but also phosphate and sulfate (Figure 3 A, B). Given that MtCEP1 is uniquely responsive to nitrogen deficiency but not phosphate or sulfate deficiency (Figure 2A), these results were unexpected. Tabata et al. 2010, did not report any change in P or S uptake in Arabidopsis in response to CEP1 application, although they did find upregulation of *AtPHT1*.*1* and *AtPHT1*.*4* in addition to NRT transporters, after 24 hours of peptide treatment. However, recent work utilizing this uptake platform to screen for genetic diversity of nutrient uptake rates in maize germplasm found that the uptakes rates of various nutrients, as well as root respiration, are generally positively correlated (Griffiths et al., 2021). This presumably reflects the need to balance uptake of different nutrients with the demand for metabolism and growth, dictated by the overall stoichiometry of elements in the plant, with faster growth requiring increased uptake of all essential nutrients and greater energy consumption. Part of this energy consumption will drive energization of cellular membranes, which in turn drives transport of various nutrients into and around cells and tissues. This may account for part of the apparent coordination in nutrient uptake observed in this and other studies. No doubt, however, full coordination requires control at many levels, including the genetic level as exemplified by changes in gene expression, as observed here.

To begin to understand how CEP peptides alter root nutrient uptake and development, we conducted RNAseq on *M. truncatula* roots three hours post treatment with the three different peptides (Figure 4). Our analysis revealed that the peptide MtCEP1D1 (1349 DEGs) had the largest effect on the Medicago transcriptome, followed by MtCEP1D2 (1278 DEGs) and AtCEP1 (617 DEGs). This is consistent with our observation that application of MtCEP1D1 on *M. truncatula* roots increases uptake of nitrate by 30% more than AtCEP1 (Figure 3). GO enrichment analysis revealed that both MtCEP1 peptide domains decreased auxin related gene expression. Repression of auxin signaling, transport and/or biosynthesis could explain the developmental changes that accompany CEP1p applications, including reduction in LR number (Figure 4B, Figure 2C). This corroborates the finding that CEP1 application represses auxin biosynthesis and alters auxin transport in Medicago roots to affect gravitropic responses in roots (Chapman et al., 2020). Moreover, application of both MtCEP1 domain encoding peptides decreased energy metabolism-related processes and sugar metabolism required for plant growth and development, consistent with the associated decrease in total root length. Enrichment of GO categories related to cell recognition (MtCEP1D1) and phosphatase activity (MtCEP1D2) are consistent with the role of MtCEP1 as a signaling peptide controlling various physiological responses. A targeted search of transporters involved in N, P, and S uptake in Medicago yielded several nitrate transporters and one sulfate transporter that were upregulated by application of CEP peptides. Increased transporter density on the root exodermis is commonly believed to enhance uptake, but other mechanisms may exist such as allelic diversity, increased assimilation to decrease internal cellular concentrations, and increased counter-ion efflux (Griffiths and York, 2020). Further functional characterization using Tnt1 insertion mutants or gene editing technologies will help to understand the contribution of specific “downstream” genes controlled by CEP1 signaling and the observed changes in root function. The absence of a clear candidate phosphate transporter that is transcriptionally regulated by the CEP peptides points to alternative mechanisms of controlling phosphate uptake under these conditions. One such possibility is the involvement of sulfate transporters in phosphate uptake, given the observation that SULTR3;5 was also shown to mediate accumulation of inorganic phosphate in rice (Yamaji et al., 2016) and our observation that SULTR genes are induced by CEP peptides in Medicago (Figure 4 C). Finally, our data also revealed novel candidate genes that may be involved in CEP1 signal perception and relay. These included several Myb-domain containing transcription factors, WRKY, GRAS domain, and ERF (AP2 ERF) transcription factors. Although a previous study overexpressing CEP1 in hairy roots of *M. truncatula* found the same family of TFs, the gene IDs were different (Imin et al., 2013) possibly due to differences in the age of plants used and the unique nature of transgenic “hairy” roots. We identified several LRR-RL kinases that were preferentially upregulated by MtCEP1D1 application (Medtr5g024420, Medtr8g068540, Medtr8g469670). This suggests that MtCEP1D1 may initiate signaling in distinct downstream pathways.

In summary, using a novel nutrient uptake analysis platform, we have found that exogenous application of specific synthetic peptides of the CEP1 family can significantly enhance nitrate uptake in Arabidopsis and Medicago by as much as 70-140% at low nutrient levels (Figure 1C). Previously, synthetic peptides have been reported to affect developmental processes . Here we show that application of a peptide can affect transcription of transporter genes and enhance nutrient uptake processes. Based on these results, SSPs show promise in horticulture, and agriculture more generally, through use in hydroponic and fertigation systems, as well as part of seed coat treatments, which would place them in close proximity to plant seedlings and roots upon germinations. Implementation of nutrient uptake enhancing SSPs in agriculture could help drive greater nutrient capture whilst minimizing nutrient losses.

## Supplementary figures

**Figure S1.**
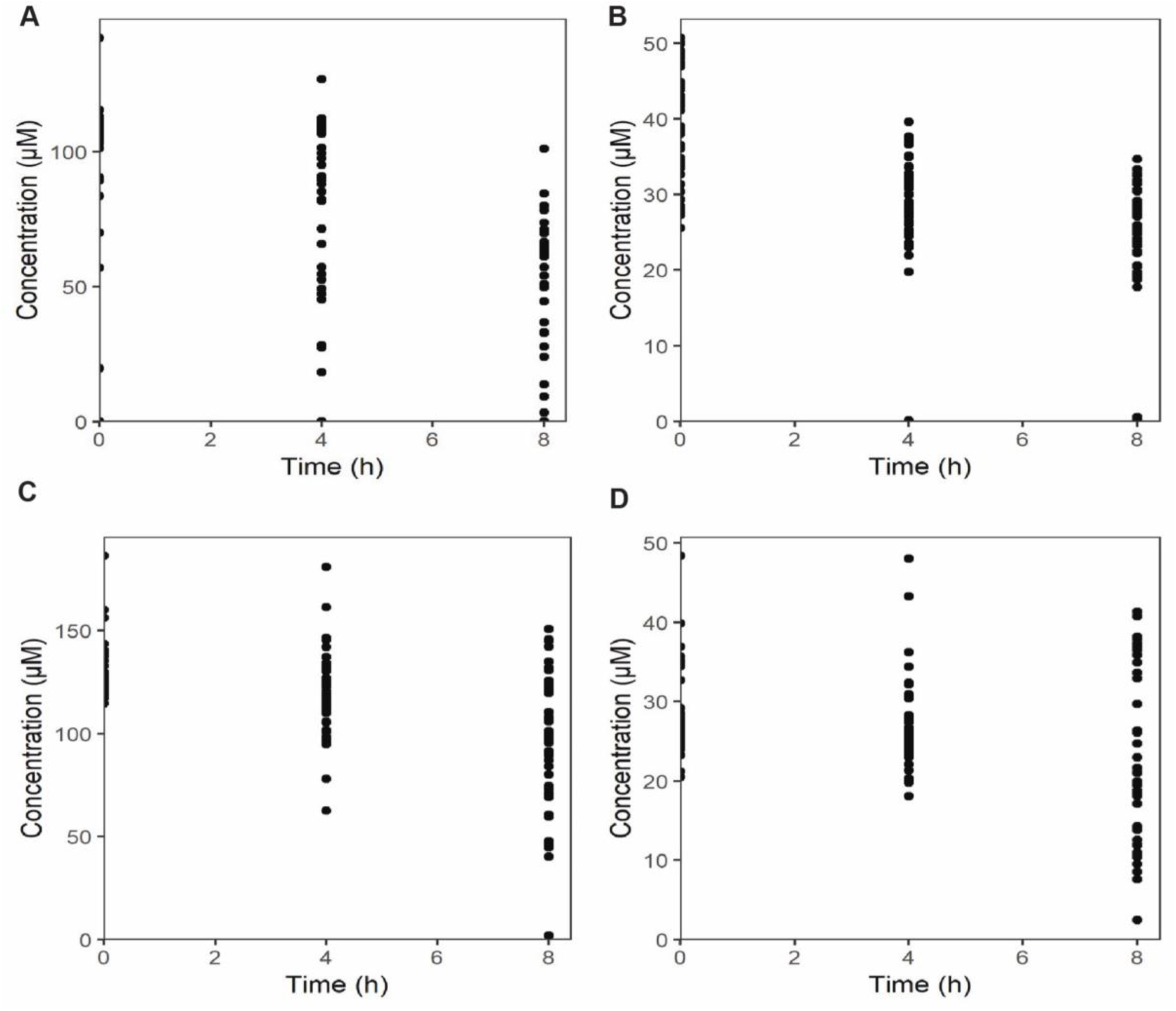
Dot plots showing nutrient uptake in 48 different plant roots treated with different peptides over 8 hours as measured by the uptake platform. Nutrient depletion plots show raw data not normalized for root length. Uptake was measured for A. Nitrate B. Phosphate C. Sulfate and D. Potassium. Each peptide and control had six replicates each.

**Figure S2.**
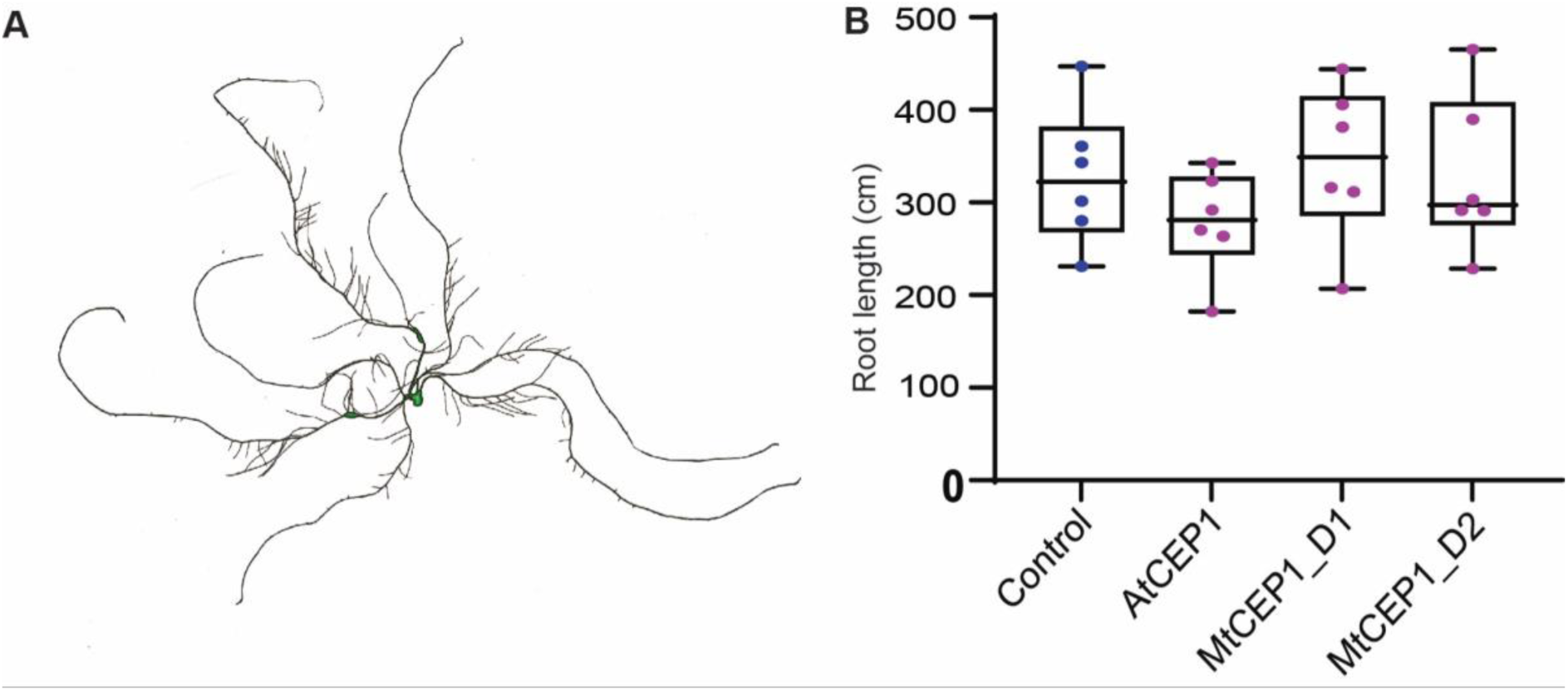
A. Rhizovision output showing representative root scan used for measuring total root length. B. Box plot showing difference in root length post treatment with peptides. No significant differences were found using a two way ANOVA followed by a Dunnett’s multiple comparison test.

**Figure S3.**
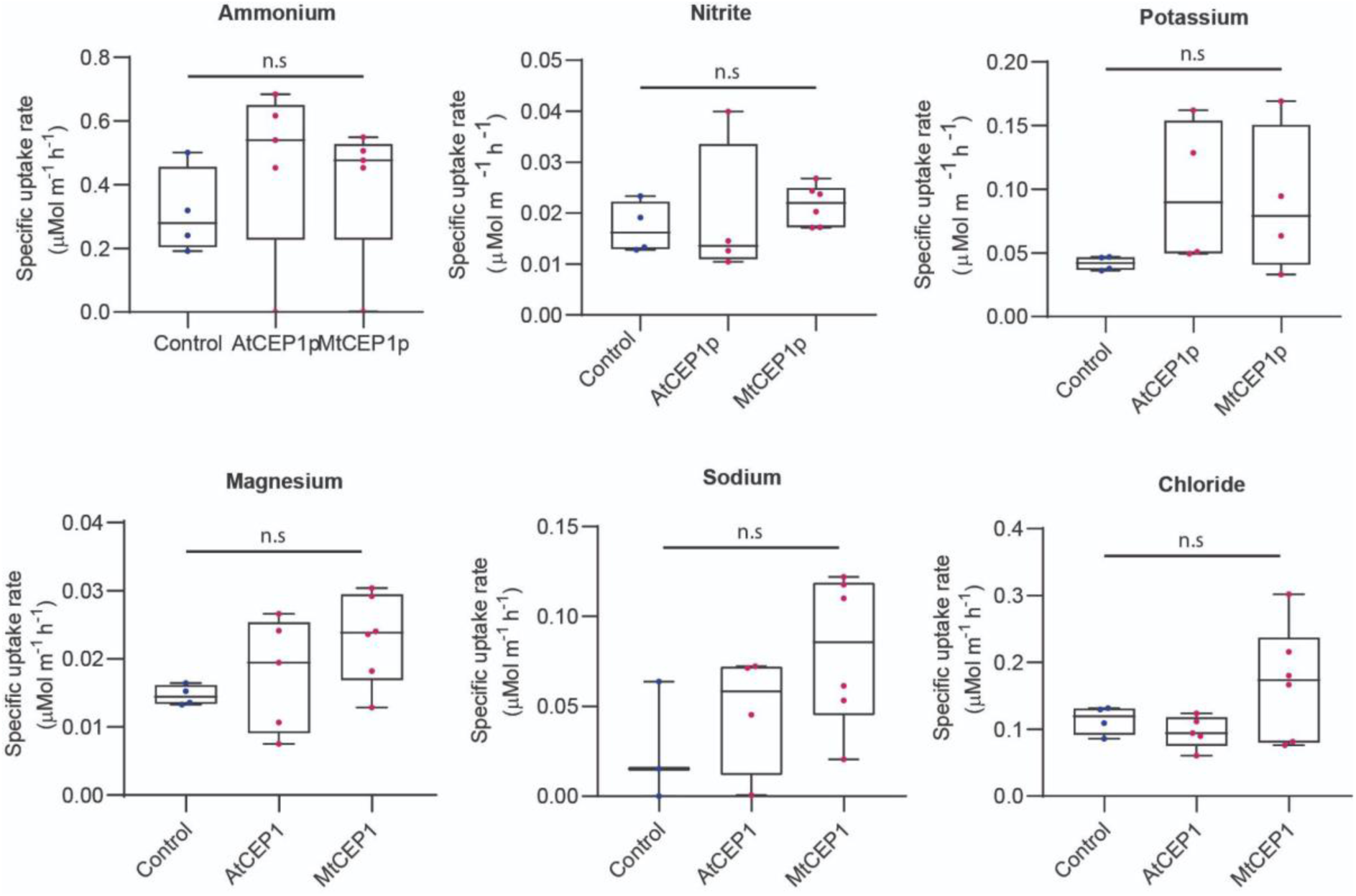
CEP1 has no significant effect on uptake of additional nutrients tested. Specific nutrient uptake rates of ammonium, nitrite, potassium, Magnesium, Sodium, Chloride as indicated in *M. truncatula* in the presence of the synthetic AtCEP1 peptide and MtCEP1D1 at a concentration of 1 μM. Student’s t-test \**p*<0.05, \*\**p*<0.01, \*\*\**p*<0.001. *n*=5-6 per treatment.

## Supplementary Tables

Supplementary Table 1: Transcript per million (TPM) counts, DESeq2 results and differentially expressed genes under MtCEP1D1, MtCEP1D2 and AtCEP1 application.

Supplementary Table 2: Gene Ontology analysis of DEGs under application of synthesized CEP1 peptide domains and unique genes in top GO terms.

Supplementary Table 3: NRT/PTR, PHT and SULTR families under application of the CEP1 peptide domains.

Supplementary Table 4: Transcription factors and kinases in top15% upregulated genes induced by CEP1 peptide domains, primer sequences and nutrient media composition.

## Acknowledgements

We would like to thank David Huhman for help with ion content measurement, Lynne Jacobs for help growing plants, and Kim Spiering for technical support. This work was funded by the National Science Foundation award #1444549 to Wolf R. Schieble and M.K Udvardi; USDA-NIFA award #2017-67007-25948 to Larry M. York; the Center for Bioenergy Innovation, a U.S. Department of Energy Bioenergy Research Center supported by the Office of Biological and Environmental Research in the DOE Office of Science; and the Lloyd Summer Noble Summer Scholar grant to S. Roy and M. Griffiths.

## Author Contributions

W.R.S, M.K.U and S.R conceptualized the peptide assays and interpreted results, L.M.Y. and M.G conceptualized and implemented the nutrient uptake measurement platform, M.G created the R scripts for data analyses, S.R. and M.G designed experiments, S.R, M.G, I.T.J, B.C, L.A, S.Z, D.J, N.D.K performed experiments and analyzed data, S.R, M.G, W.R.S, M.K.U, L.M.Y, wrote this manuscript with input from all authors.

